# Manual lymph drainage massage of the head and neck improves cognition and reduces pathological biomarkers in the *5x-FAD* mouse model of Alzheimers disease

**DOI:** 10.1101/2025.08.08.669361

**Authors:** Mitchell J. Bartlett, Robert P Erickson, Jennifer Frye, Kristian P. Doyle, Paulo W. Pires, Marlys H. Witte

## Abstract

Alzheimer’s disease (AD) affects 6.9 million people over the age of 65 in the US and is expected to double by 2060. While FDA approved immunotherapies slow cognitive decline in some individuals with AD, they do not improve cognition, are costly, and have significant side-effects. Therefore, new targets, approaches, and treatments for AD are a necessity. There are no FDA approved therapies for AD that target the brain’s lymphatic system. It is well established that the toxic protein, amyloid-beta (Aβ), accumulates in the AD brain. Recent studies have shown that Aβ is cleared via interstitial fluid and cerebrospinal fluid through a pathway involving the glymphatic system—meningeal lymphatic vessels—leading to deep and superficial cervical lymphatic vessels and nodes. Therefore, any blockage along this route can cause inefficient drainage and result in pathological buildup of Aβ, which can lead to AD.

Here, we propose a new approach to treating AD by manual lymph drainage (MLD), which is a light skin massage traditionally used to reduce fluid accumulation in lymphedema. This therapy has also been demonstrated to be safe in individuals with AD, but its effects on cognition and biomarkers of AD has never been investigated. In this study we demonstrate that repeated MLD of the head and neck, including the superficial cervical lymphatic vessels (scLVs), improves cognitive function in AD as measured in both the Y-maze and nest-building tests. We also show that this coincides with a reduction in plasma levels of neurofilament light chain (NfL), a non-specific biomarker for neuronal cell death and axonal damage. MLD was also shown to reduce Aβ in the hippocampus of these mice. Combined, this data provides compelling proof-of-principle evidence for the potential of MLD as a standalone or adjunct therapy in the treatment of AD.

## 1 INTRODUCTION

Alzheimer’s disease (AD) affects approximately 6.7 million Americans over the age of 65, a number expected to increase to 13.8 million by 2060.[1] Globally, AD has a much greater impact. Considering the entire AD continuum (preclinical AD, prodromal AD, and AD dementia), there are currently an estimated 416 million individuals affected and at risk. [2] However, current treatment options are limited. Therefore, there is a critically unmet need for novel therapeutic targets and innovative treatments, specifically those that aid in AD prevention and are accessible to all individuals.

Manual lymph drainage (MLD) is a non-invasive technique involving light manipulation of the skin, that increases contractility, lymph flow, and protein clearance.[3–5] It is a well-established therapy and is used in the first phase of complex decongestive therapy (CDT) for individuals with peripheral lymphedema.[6] However, MLD has also been demonstrated to alleviate intracranial pressure,[7] cerebral edema and head pain in post-traumatic brain injury,[8] while also reducing dementia associated agitation,[9] and increasing social contact which protects against dementia.[10] MLD is also inexpensive, and can be self-administered or performed by a trained family member or caregiver, making it easily accessible.

The brain’s lymphatic system, which includes the interconnected glymphatic, meningeal lymphatic, and cervical lymphatic pathways, plays a key role in clearing cerebrospinal fluid (CSF) and interstitial fluid (ISF), and provides a compelling therapeutic target.[11–14] This is supported by preclinical rodent models and human studies, both of which have implicated that impaired function of the glymphatic [11,15–24] and the meningeal lymphatic vessels [13,25–27] plays a pivotal role in the removal of CSF from the brain. However, interplay between the cervical lymphatic vessels (cLVs) and AD is much less clear. Preclinical models have demonstrated that deep cervical lymphatic vessel (dcLV) ligation worsens pathology in a mouse model of AD. However, it is unknown if this vessel is affected in any established preclinical transgenic model or clinical AD. There is even less known about the impact of superficial cervical lymphatic vessels (scLV) impairment on AD. However, there is evidence that CSF, containing Aβ, exits the brain through the scLVs as demonstrated by the accumulation of Aβ in the superficial cervical lymph nodes of both mice [28] and humans.[29] This provides further evidence for these vessels as a potential therapeutic target.

This study is the first of its kind to demonstrate that MLD improves spatial working memory and activities of daily living, as measured by spontaneous alternation and the nest building test, respectively. Last, we showed that MLD also improved biomarkers of AD, significantly reducing both plasma levels of neurofilament light chain (NfL) and the Aβ_42/40_ ratio in the hippocampus of *5x-FAD* mice.

## 2 METHODS

### 2.1 Animals

This study used 5-month-old female *5x-FAD* mice. This is a model of early-onset AD that combines five different mutations in two genes associated with familial AD. Three of the mutations are in the gene that codes human amyloid precursor protein (APP): Swedish (K670N, M671L), Florida (I716V), and London (V717I). The other two mutations are inserted in the gene of human presenilin, M146L and L286V. Transgene expression is controlled by the neuronal-specific *Thy1* promoter.[30] *5x-FAD* transgenic mice (Strain No. 34840-JAX) were purchased from The Jackson Laboratory (Bar Harbor, ME) and are actively maintained in our breeding colony at The University of Arizona College of Medicine (UACOM). All mice were bred on a C57BL/6J background, and wild-type (WT) littermates were used as controls. Mice were co-housed, 4-5 mice per cage (20.7 × 31.6 × 21.5 cm), within isolated Innovive racks in a temperature (25°C ± 1°C) and humidity-controlled room on a 12-hour light/dark cycle. Food and water were available ad libitum.

All animal procedures in this study were approved by the Institutional Animal Care and Use Committee of the University of Arizona College of Medicine (IACUC protocols 15-053 and 18-473) and are in accordance with the National Institutes of Health’s Guide for the Care and Use of Laboratory Animals, 8th edition. All animal experiments are reported in compliance with Animal Research: Reporting of In Vivo Experiments (ARRIVE) guidelines 2.0.

### 2.2 Experimental Design

*5x-FAD* mice underwent MLD or sham treatment for 15-minutes, twice per day, for 67 days. MLD and sham treatment continued during behavioral testing phase (Days 61-67), which was immediately followed by extraction of brain tissue for post-mortem biochemical analysis.

### 2.3 Manual Lymph Drainage

Manual lymph drainage (MLD) is a gentle, light pressure, technique that involves stretching the skin to stimulate lymphatic vessel contraction, promoting lymph movement towards the lymph nodes. In this study, we adapted the Vodder method of MLD for mice.[31] Mice were brought to a dedicated treatment room daily for the duration of the study (*see 2.2 Experimental* Design), one hour before treatment. To not disrupt their circadian rhythms, red lights were affixed, and the temperature and humidity were adjusted to match their colony and behavioral testing rooms. A single experimenter provided all treatment sessions, which were applied twice per day, at four- and eight-hours hours after the onset of their dark cycle. Each treatment session lasted approximately 20-minutes. In the three-days prior to beginning the experiment, all the mice underwent sham treatment to acclimate them to the experimenter and treatment room. Sham treatment consisted of holding each mouse at the scruff for 20-minutes with no other physical touch. For the WT and *5x-FAD* mice sham treatment continued for the remainder of the experiment. MLD treated mice (MLD+*5x-FAD*) were held at the scruff and then underwent MLD. All locations of MLD treatment were performed caudally to cranially, anatomically. MLD treatment was also divided into two phases. The goal of Phase One was to prime and drain the lymph nodes for receiving lymph fluid. This was performed using gentle circular motions with the thumb and index finger, providing bilateral stimulation, for 30-seconds to drain the following lymph nodes in this order: abdominal, axillary, supraclavicular, superficial cervical, and preauricular lymph nodes. Phase Two involved moving the lymph, which consisted of gently and rhythmically stretching the skin in a downward motion and then allowing it to return to its natural position. This was performed 10 times each on the skin covering the following anatomical locations in this order: acromion, collarbone, side of the neck, superficial lymphatic vessels (throat), mandible, back of neck, base of the skull, preauricular, postauricular, naso orbital, retroorbital, and skull. Phase Two was then repeated twice, for a total of three cycles.

### 2.4 Behavioral Tests

Behavioral tests were performed in our dedicated Mouse Neurobehavior Suite. To acclimate, mice were transported from the adjacent treatment room one-hour before testing. All behavioral tests were performed by the same experimenter, during the animals’ dark cycle, under red light to avoid disrupting their circadian rhythms. Additionally, as treatment was ongoing during the testing phase, each test was performed two-hours after the first sham or MLD treatment and completed at least 1-hour before the second treatment. A 24-hour rest period was allowed between tests, which were performed in the following order: spontaneous alternation (Y-Maze) and nest building test. All testing arenas and apparatuses were cleaned in between animals with a 70% ethanol solution and allowed to dry before subsequent tests. Tests were recorded with a camera phone (iPhone 8; Apple, Cupertino, CA) and then analyzed, when noted, with video tracking software (ANY-maze, Version 7.49; Stoelting Co., Wood Dale, IL).

#### 2.4.1 Spontaneous Alternation

Using methods established in the *5x-FAD* mouse model, spontaneous alternation was used to assess spatial working memory and exploratory behavior. [32,33] Spontaneous alternation was measured with an opaque Y-maze (in-house fabrication; arms were 5 x 40 x 10 centimeters). Mice were placed in the center of the maze and the percentage of spontaneous alternations and total number of entries into each arm was measured over a 5-minute period. The percentage of spontaneous alternations was calculated by the formula: [(number of alternations)/(total number of arm entries – 2) x 100%]. An alternation was defined when a mouse entered each of the three independent arms consecutively without entering a previously visited arm.

#### 2.4.2 Nest Building Test

The nest building test was used to measure executive function or the ability to perform activities related to daily living.[34,35] Each mouse was moved from co-housing to an individual clean cage. Mice were allowed to acclimate to individual housing for 24-hours to reduce anxiety and stress following separation from their littermates. After the acclimation period, but one-hour before the next 12-hour dark cycle, two pre-weighed (5.0 grams) cotton nestlets were placed in each cage to allow the mouse to build nests. Nests were assessed 24-hours later using a rating scale of 1–5 as established in our laboratories.[35] Additionally, untorn nestlet pieces, defined as those greater than 0.1 grams or approximately 4 percent of a nestlet, were weighed and subtracted from original weight to quantify nest building.[34]

### 2.5 Plasma & Tissue Collection

Mice were euthanized following behavioral testing on Day 67. They were deeply euthanized with isoflurane and a transcardial catheter was placed. Blood was drawn through the catheter and placed in K3EDTA treated tubes (CT-M500-K3EDTA; Sarstedt AG & Co. KG, Nümbrecht, Germany). Samples were gently inverted 10-times and then centrifuged at 4 °C for 10-minutes at 2,000 x g. Plasma was aliquoted into sterile tubes, flash frozen in liquid nitrogen and stored at −80 °C. Following blood collection, the catheter was then used to perfuse each mouse with 12 mL of ice-cold phosphate buffered saline (PBS). The brain was extracted, and regions of interest, including the hippocampus and prefrontal cortex, were dissected, flash frozen in liquid nitrogen, and stored at −80 °C for analysis.

### 2.6 Sample Preparation, Normalization, and Immunoassays

#### 2.6.1 Normalization of Plasma and Brain Tissue Samples

A subset of samples from the cohort of animals used for behavioral testing were randomly selected for use in the biochemical assays described below. One plasma sample from each treatment group was removed due to hemolysis. Each plasma sample was diluted 1:8 with the kit provided diluent. Brain tissue lysate was prepared as previously published with a few modifications.[36] After homogenization, the tissue lysate was centrifuged at 4 °C for 1-hour at 17,000 x g to remove the insoluble fraction. Total protein of the soluble fraction was determined using a Pierce^TM^ BCA Protein Assay Kit (Cat. No. 23225; ThermoFisher Scientific, Waltham, MA) and quantified on a FlexStation 3 microplate plate reader (Molecular Devices, San Jose, CA). Lysate from each hippocampal and prefrontal cortex sample was then normalized with our previously described radioimmunoprecipitation assay buffer to 1.0 mg/mL.

#### 2.6.2 Neurofilament Light Chain Assay

Neurofilament light chain (NfL) was quantified using a Simoa® (Single Molecule Array) NF Light v2 Advantage Assay (Cat. No. 104073; Quanterix®, Billerica, MA). This assay has been established in our laboratory[37] and was carried out according to the manufacturer’s instructions.

#### 2.6.3 Amyloid Beta Assay

Amyloid-beta (Aβ) was quantified using a MILLIPLEX® human amyloid beta and tau magnetic bead panel-multiplex assay (Cat. No. HNABTMAG-68K; MilliporeSigma, Burlington, MA). The assay was carried out according to the manufacturer’s instructions.

### 2.7 Statistical Analysis

Statistical analyses were performed using Prism (Version 10.0; GraphPad Software, La Jolla, CA). The null hypothesis was rejected when *p* < 0.05. All values are represented as mean ± SEM. Behavior and biochemical data were analyzed by one-way ANOVA, with a Fisher’s LSD *post hoc* test. The qualitative assessment of the nest building test was analyzed with a non-parametric Kruskall-Wallis test, followed by an uncorrected Dunn *post hoc* test.

## 3 Results

### 3.1 Manual lymph drainage improves cognitive function in *5x-FAD* mice

The impact of manual lymph drainage on cognitive function was first investigated in the Y-maze test (Figure 1). WT mice had a percent spontaneous alternation (mean ± SEM) of 64.1 ± 4.1%, which was significantly decreased by 28% in *5x-FAD* mice to 46.4 ± 6.5% (WT vs. *5xFAD*: *t* = 2.09, df = 21, *p* < .01, *n* = 6-12 per group). The cognitive impairment in the *5x-FAD* mice was reversed by MLD. Specifically, MLD+*5x-FAD* mice not only significantly improved by 45% to 67.3 ± 9.3% compared to sham treated *5x-FAD* mice, but they also performed at the same level as WT controls (MLD+*5x-FAD* vs. *5x-FAD*: *t* = 2.13, df = 21, *p* < 0.01, *n* = 6 per group; MLD+*5x-FAD* vs. WT: *t* = 0.36, df = 21, *p* = 0.72, *n* = 6-12 per group). Importantly, the number of arm entries (mean ± SEM) between WT (17.8 ± 0.9), *5x-FAD* (21.5 ± 2.2), and MLD+*5x-FAD* (18.8 ± 1.9) mice was not significantly different (WT vs. *5xFAD*: *t* = 1.76, df = 21, *p* = 0.09, *n* = 6-12 per group; MLD+*5x-FAD* vs. *5x-FAD*: *t* = 1.09, df = 21, *p* = 0.29, *n* = 6 per group; MLD+*5x-FAD* vs. WT: *t* = 0.51, df = 21, *p* = 0.62, *n* = 6-12 per group).

**Figure 1.**
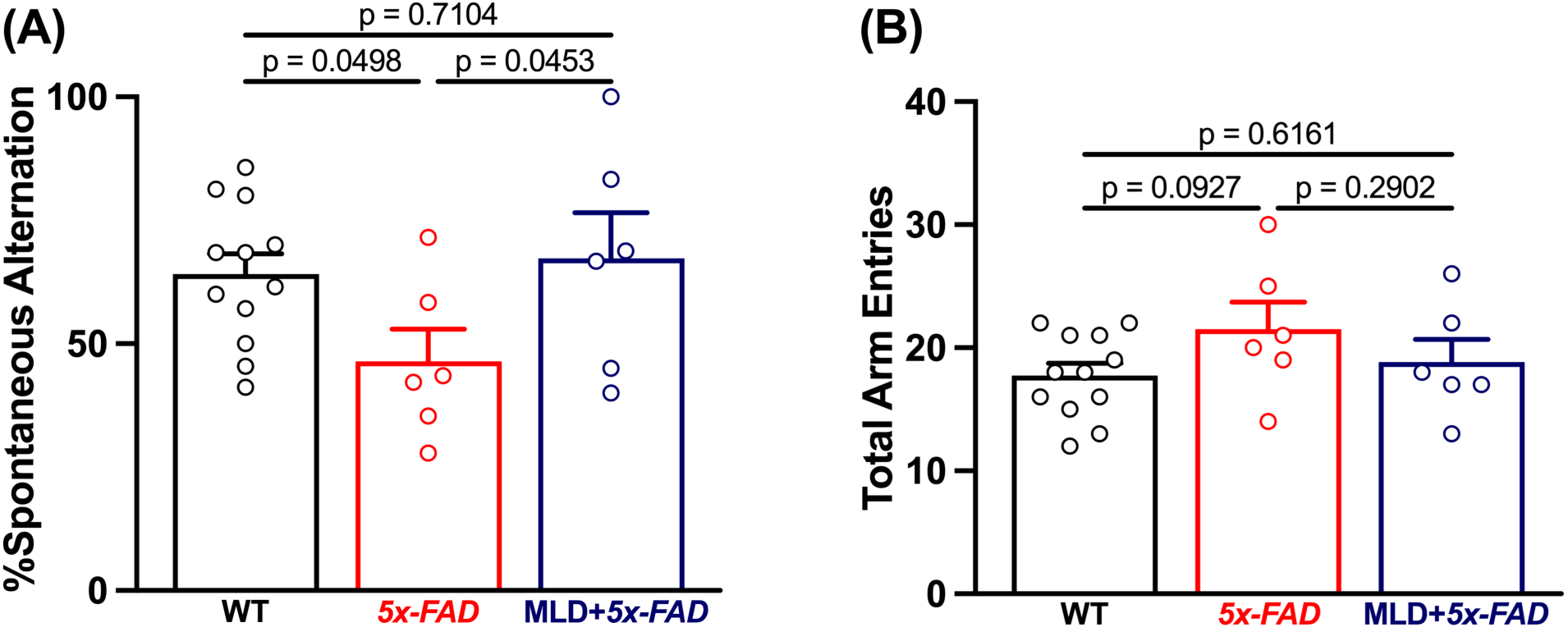
Manual lymph drainage (MLD) improves spatial working memory in *5x-FAD* mice. (A) Summary bar graph showing the effect of MLD on spontaneous alternation in the Y-Maze. *5x-FAD* mice have impaired spatial working memory, which is reversed to WT levels by MLD. (B) Summary bar graph of the total number of arm entries by each group during the Y-maze test. There were no significant differences between groups, demonstrating that the impaired memory in *5x-FAD* mice and the reversal by MLD in (A) was not due to a difference in exploratory behavior. Each individual data point represents one mouse. All data are represented as mean ± SEM. Statistical analysis: one-way ANOVA, with a Fisher’s LSD *post hoc* test.

To further evaluate cognitive function, the mice were challenged with the nest building test (Figure 2). Like the Y-maze, *5x-FAD* mice showed a significant impairment in both qualitative (Figure 2A,B) and quantitative (Figure 2C) nest building measures as compared to WT mice. MLD significantly improved these measures. To assess the completeness or quality of the nests, each was assigned a nest-rating score (mean ± SEM) based on shape and structure (Figure 2A). WT mice had a qualitative nest-rating score of 3.6 ± 0.3, which was significantly reduced in *5x-FAD* mice by 71% to 1.0 ± 0.02 (WT vs. *5xFAD*: *Z* = 3.55, *p* < 0.001, *n* = 6-9 per group). MLD+*5xFAD* nests had a nest-rating score of 2.9 ± 0.3, a significant improvement of 183% compared to sham treated *5x-FAD* mice, and not significantly different than WT mice (MLD+*5x-FAD* vs. *5x-FAD*: *Z* = 2.54, *p* < 0.01, *n* = 6 per group; MLD+*5x-FAD* vs. WT: *Z* = 0.77, *p* = 0.44, *n* = 6-9 per group). In addition to the qualitative nest-rating score, the amount of the nestlets torn (mean ± SEM grams) by each mouse was also quantified. WT mice tore 4.0 ± 0.2 grams of their nestlets. *5x-FAD* mice tore 1.5 ± 0.4 grams, a significant reduction of 63% compared to WT mice (WT vs. *5x-FAD*: *t* = 4.39, df = 18, *p* < 0.001, *n* = 6-9 per group). MLD+*5x-FAD* tore 2.9 ± 0.6 grams, a significant 89% improvement compared to sham treated *5x-FAD* mice (MLD+*5x-FAD* vs. *5x-FAD*: *t* = 2.13, df = 18, *p* < 0.05, *n* = 6 per group). There was also no significant difference between the treated MLD+*5x-FAD* mice and WT mice (MLD+*5x-FAD* vs. WT: *t* = 2.06, df = 18, *p* = 0.054, *n* = 6-9).

**Figure 2.**
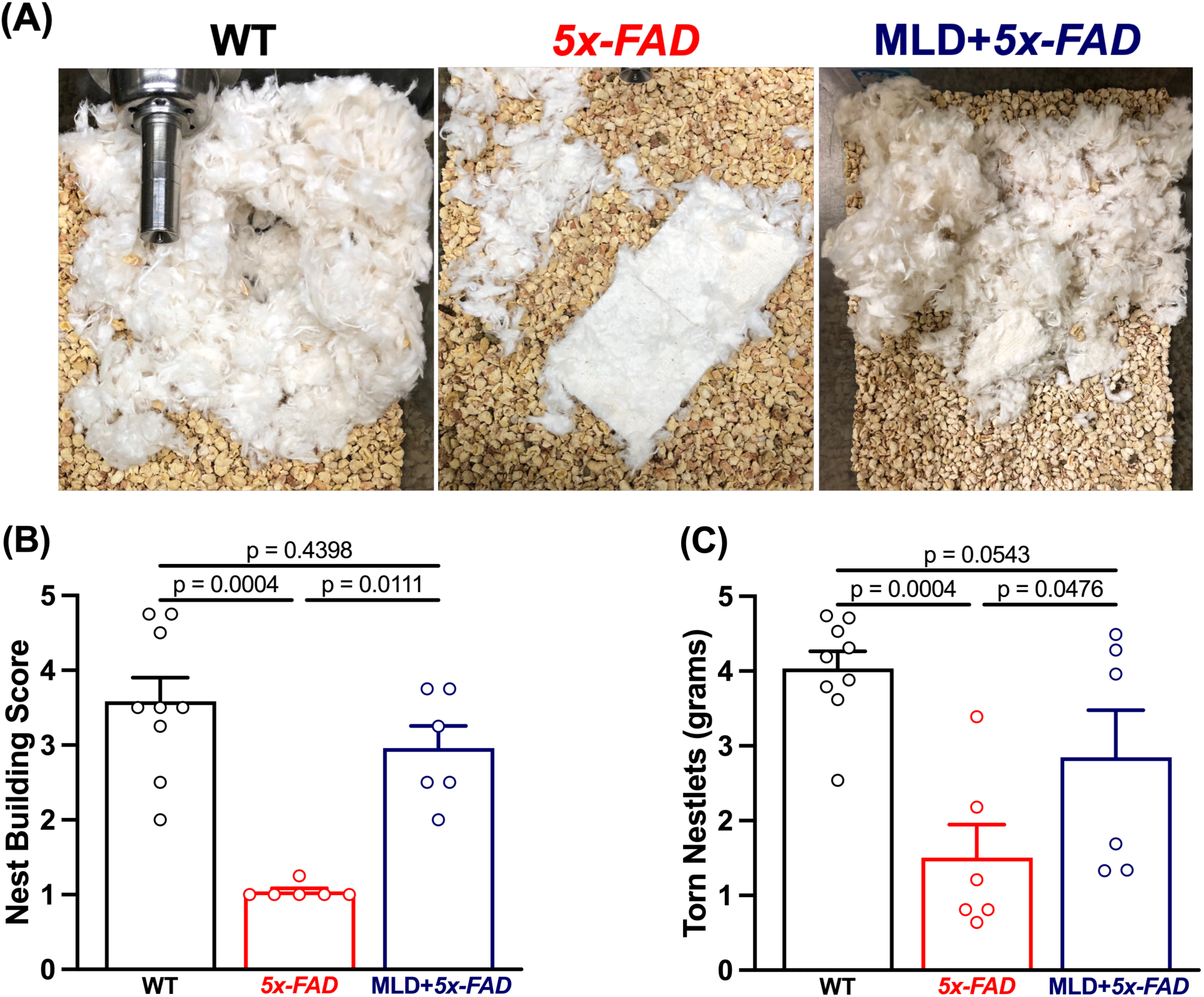
Manual lymph drainage (MLD) improves nest building in *5x-FAD* mice. (A) Representative images of nests made by WT (left image), sham treated *5x-FAD* mice (middle), and MLD treated *5x-FAD* mice (right). Note the incomplete nests in the *5x-FAD* mice (middle) as compared to the partially restored nests in the MLD treated *5x-FAD* mice (right). (B) Summary bar graph of the nest building scores show that female *5x-FAD* mice build significantly incomplete nests and that these are significantly improved in *5x-FAD* mice treated with MLD. Each individual data point represents one mouse. All data are represented as mean ± SEM. Statistical analysis: (B) Qualitative assessment of the nest building scores was analyzed with a non-parametric Kruskall-Wallis test, followed by an uncorrected Dunn *post hoc* test. (C) Quantitative assessment of one-way ANOVA, with a Fisher’s LSD *post hoc* test.

**Figure 3.**
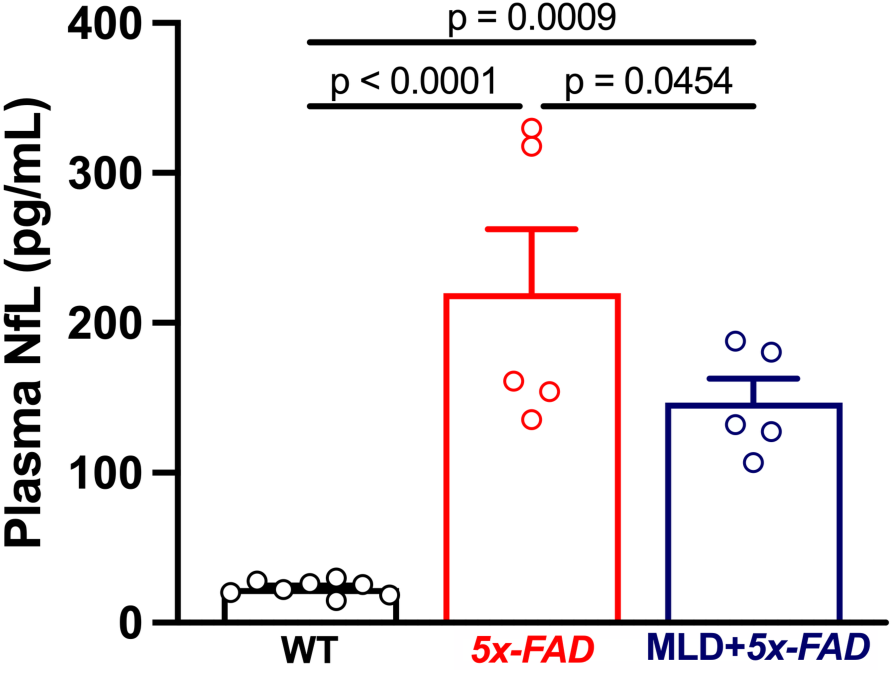
Manual lymph drainage (MLD) reduces plasma Neurofilament Light Chain (NfL) in *5x-FAD* mice. Summary bar graph showing the effect of MLD on plasma NfL levels. Both sham treated and MLD treated *5x-FAD* mice have significantly elevated plasma NfL compared to WT mice. However, MLD significantly reduced plasma NfL compared to sham treated *5x-FAD* mice suggesting a therapeutic effect. Each individual data point represents one plasma sample from a single mouse. All data are represented as mean ± SEM. Statistical analysis: one-way ANOVA, with a Fisher’s LSD *post hoc* test.

**Figure 4.**
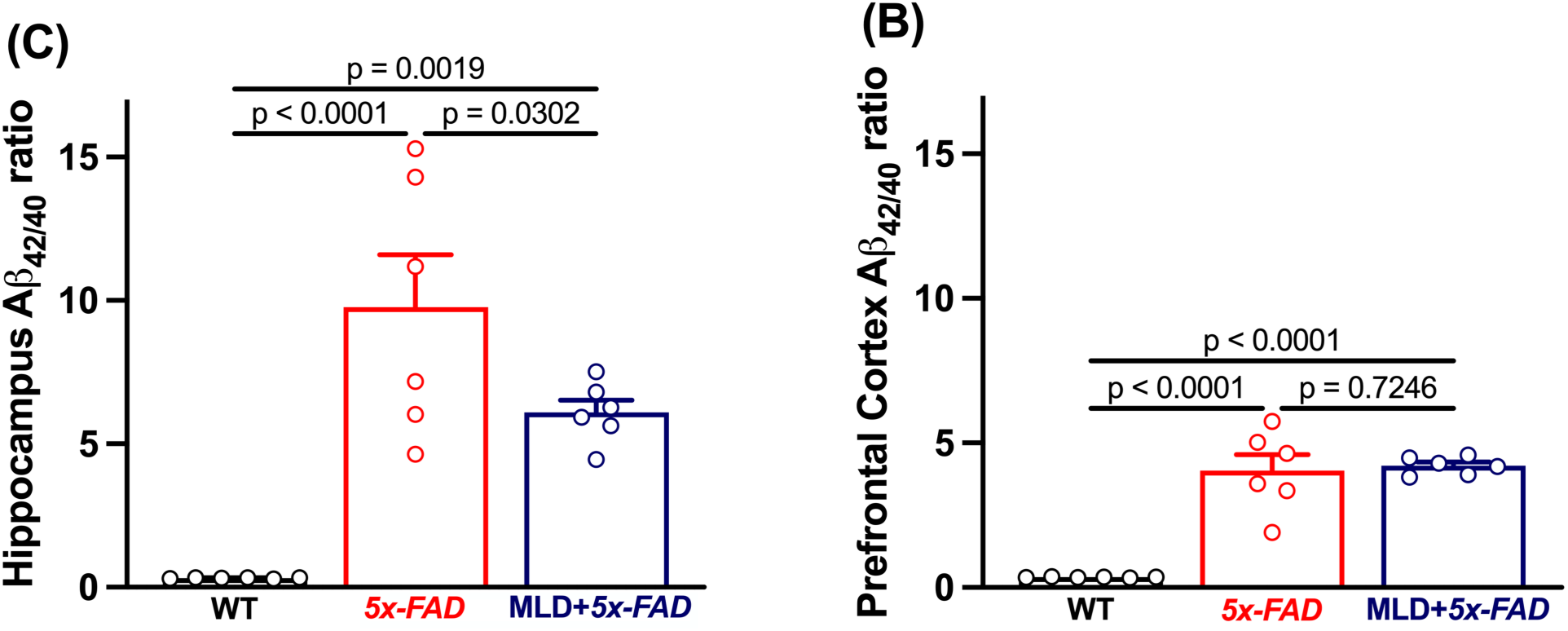
Manual lymph drainage (MLD) reduces amyloid-beta (Aβ) in the hippocampus, but not prefrontal cortex. (A) Summary bar graph showing the effect of MLD on the Aβ_42/40_ ratio in the hippocampus. Aβ is significantly increased in the hippocampus of sham treated and MLD treated *5x-FAD* mice compared to WT mice. MLD significantly reduced the hippocampal Aβ_42/40_ ratio compared to sham treated *5x-FAD* mice suggesting a therapeutic effect. (B) Summary bar graph showing the effect of MLD on the Aβ_42/40_ ratio in the prefrontal cortex. Aβ is significantly increased in the prefrontal cortex of both sham treated and MLD treated *5x-FAD* mice compared to WT mice. There was no significant effect of MLD on the Aβ_42/40_ ratio in the prefrontal cortex. Each individual data point represents either one hippocampal or prefrontal cortex sample from a single mouse. All data are represented as mean ± SEM. Statistical analysis: one-way ANOVA, with a Fisher’s LSD *post hoc* test.

### 3.2 Manual lymph drainage reduces plasma Neurofilament Light Chain in *5xFAD* mice

Following behavioral testing, we investigated the effect MLD had on pathological biomarkers related to AD. First, we looked at the effect of MLD on plasma neurofilament light chain (NfL), a non-specific biomarker that has been used to monitor neurodegeneration in preclinical and clinical AD.[38] As expected, NfL plasma levels (mean ± SEM pg/mL) in WT mice were very low, 22.8 ± 1.8, whereas plasma NfL was significantly increased in *5x-FAD* mice to 220 ± 42.8 pg/mL, an increase of 865% (WT vs. *5xFAD*: *t* = 6.55, df = 15, *p* < 0.0001, *n* = 5-8 per group). Following treatment, plasma NfL significantly decreased by 33.18% to 147 ± 15.8 pg/mL in the treated MLD+*5xFAD* mice compared to untreated *5xFAD* mice (MLD+*5xFAD* vs. *5xFAD*: *t* = 2.18, df = 15, *p* < 0.05, *n* = 5 per group). However, treatment did not reduce plasma NfL in MLD+*5xFAD* mice to WT levels. The MLD+*5xFAD* mice still showed significantly greater plasma NfL, specifically a 545% increase, compared to WT mice (MLD+*5xFAD* vs. WT: *t* = 4.13, df = 15, *p* < 0.001, *n* = 5-8 per group).

### 3.3 Manual lymph drainage reduces Aβ in the hippocampus but not the prefrontal cortex

To better elucidate the impact of MLD on AD, we analyzed one of its hallmark pathological biomarkers, the Aβ_42/40_ ratio (mean ± SEM), in both the hippocampus and prefrontal cortex. This biomarker has been shown to be higher in the *5x-FAD* model and increase in the brain with age.[30,39] As expected, the WT mice had a very low Aβ_42/40_ ratio, 0.32 ± 0.01, in the hippocampus. The Aβ_42/40_ ratio of sham treated *5x-FAD* mice was 9.8 ± 1.8, which was a significant increase of 2,963% from WT mice (WT vs. *5xFAD*: *t* = 6.16, df = 15, *p* < 0.0001, *n* = 6 per group). MLD significantly reduced the hippocampal Aβ_42/40_ ratio in MLD+*5x-FAD* mice to 6.10± 0.4, a 38% decrease compared to sham treated *5x-FAD* mice (MLD+*5xFAD* vs. *5xFAD*: *t* = 2.39, df = 15, *p* < 0.05, *n* = 6 per group). Despite the improvement MLD provided, as compared to *5x-FAD* mice, the hippocampal Aβ_42/40_ ratio in MLD+*5x-*FAD mice remained significantly greater than WT mice by 1,812% (MLD+*5xFAD* vs. WT: *t* = 3.77, df = 15, *p* < 0.01, *n* = 6 per group).

In the prefrontal cortex, the Aβ_42/40_ ratio in WT mice was 0.35 ± 0.4. The Aβ_42/40_ ratio of sham treated *5x-FAD* mice was 4.0 ± 0.6, and for treated MLD+*5x-FAD* mice it was 4.2 ± 0.1. Respectively, these were a significant increase of 1,041% and 1,089% compared to the Aβ_42/40_ ratio of WT mice (WT vs. *5xFAD*: *t* = 7.85, df = 15, *p* < 0.0001, *n* = 6 per group; WT vs. MLD+*5xFAD*: *t* = 8.20, df = 15, *p* < 0.0001, *n* = 6 per group). MLD did not reduce Aβ_42/40_ in the prefrontal cortex. There was no difference between the *5x-FAD* and MLD+*5x-FAD* mice (MLD+*5xFAD* vs. *5x-FAD*: *t* = 0.36, df = 15, *p* = 0.73, *n* = 6 per group).

## 4 Discussion

The evidence for the importance of amyloid-beta (Aβ) clearance by the brain’s lymphatic vessels has been well-established. [11,12,27,28,40] However, our study is the first of its kind to investigate the use of a non-invasive manipulation of the head and neck lymphatics, by manual lymph drainage (MLD), as a potential therapy for Alzheimer’s disease (AD). This study has many strengths, including demonstrating that MLD can be adapted to and performed repeatedly on awake mice, without anesthesia, for more than 2-months. This is important as non-invasive translational therapies proposed in the preclinical literature have regularly been performed in anesthetized mice.[41,42] Additionally, all treatments were performed over an extended period to mirror what might be done clinically or potentially at home by self-administration or via a caretaker.

An additional strength was the use of the *5x-FAD* mouse model, which is well-established and characterized by early onset and a rapid acceleration of disease pathology, while also demonstrating several hallmarks of AD that mirror those seen in humans.[30] MLD was initiated at 5-months of age to assess the impact of treatment after the onset of Aβ accumulation and cognitive impairment, which have been shown to occur in female *5x-FAD* mice as early as 1.5- and 3.5-months of age, respectively.[30,43] This experimental design provided a more realistic translational approach, as therapies are most often initiated after the onset of symptoms and a confirmed diagnosis. However, because MLD is non-invasive and without side-effects, a prevention study whereby MLD is administered before the onset of symptoms would be therapeutically and translationally relevant.

For the MLD study, we only used females, which in this model show an earlier onset of Aβ accumulation and cognitive impairment.[30] This is relevant as following advanced age, genetics and female sex are two major risk factors for an AD diagnosis; women comprise nearly two-thirds of the individuals living with AD.[1,44] This is supported by data showing that healthy middle-aged women, without an AD diagnosis, have significantly more Aβ than men[45] Two additional studies have also demonstrated a strong trend for greater Aβ burden in women.[46,47]

Behavioral studies were used to assess the effect of MLD on cognitive function. Specifically, spontaneous alternation in the Y-maze and both qualitative (nest-rating score) and quantitative measures (weight of torn nestlets) in the nest-building test were analyzed. After validating the AD phenotype of 7-month-old, female *5x-FAD* mice in both tests, we then demonstrated that MLD significantly improved performance compared to sham treated *5x-FAD* mice and brought the MLD+*5x-FAD* back to WT performance levels. It is also important to note, that while both tests assess memory and are hippocampal/prefrontal cortex-dependent, nest-building also measures motor skills, general well-being, and is the closest behavioral parallel to activities of daily living (ADLs), which are affected at multiple stages along the AD continuum.[48] Therefore, in future studies it will be important to consider other behavioral readouts, such as motor function, anxiety, and other measures of memory or cognitive function.

The biochemical data showed that neurofilament light chain (NfL) was significantly increased in the sham treated *5x-FAD* mice and this was partially reversed by MLD. NfL is one type of neurofilament that forms key protein scaffoldings in nerves and is increased in the blood and CSF in response to neuronal and axonal injury. Therefore, it has been proposed as a potentially important and reliable biomarker for neuronal injury, that also correlates with the severity of neurodegeneration.[49] Likewise, we saw similar results in our measurement of the Aβ_42/40_ ratio, a major hallmark of AD, which has been shown to increase with age in the *5x-FAD* model. Here, MLD significantly reduced the Aβ_42/40_ ratio in the hippocampus compared to sham treated *5x-FAD* mice. This biomarker data, combined with our behavioral results, strongly suggests that MLD may improve lymph flow, which is known to be reduced with age[50] and worsen AD when impaired,[40] without impairing the intrinsic contractility of the lymphatic vessels. This hypothesis is supported by a recent study from, Jin, et al., in which they designed a mechanical device to massage facial and cervical lymphatics to increase lymph flow. [42] Their group also demonstrated that CSF drainage via the scLVs is impaired with aging, which is the greatest risk factor for AD, and that this impairment can be temporarily reversed in anesthetized mice with low-magnitude mechanical stimulation of the scLVs.[42] These results lend further proof-of-principle evidence for our investigation of the impact of AD on scLV function and significantly strengthen our enthusiasm for MLD as a treatment for AD.

Currently, there are a few translational therapies that have been proposed in the preclinical literature demonstrating improvements in glymphatic and meningeal lymphatic function. These approaches include cranial bone maneuvers[51], focused ultrasound[41], growth factors[13,27,52], and pharmacological interventions [53–58]. While all show translational potential, to increase CSF outflow each requires either an invasive technique or systemic administration, which in contrast to MLD exposes future patients to potential side-effects. While several elegant studies, using MRI and fluorescent tracers, have further delineated the pathways by which the brain clears CSF, the exact mechanisms by which this occurs is still unclear. Therefore, we acknowledge that more information is needed related to the role that the lymphatics play in CSF clearance in humans to assure MLD’s success in a clinical trial. Similarly, we do not know the exact mechanism by which MLD works, which was not investigated in this initial study but will be important for furthering the understanding of MLD and its therapeutic potential. One hypothesis is that MLD works by stimulating mechanosensitive non-selective cationic ion channels, like Piezo1. This mechanoreceptor is in the endothelium, responds to stretch, and when activated generates a favorable electrochemical gradient that induces a contractile wave through the lymphangion, increasing fractional pump flow.[59]

In conclusion, there is no approved therapy that directly targets the brain’s lymphatic system for the treatment of AD, or any neurodegenerative disease. Moreover, the treatment options that do exist for AD are few, and those that do exist are often expensive, come with significant side-effects, and most importantly are not disease-modifying. Therefore, new innovative ideas are needed. MLD has previously been used for the treatment of lymphedema but has also been shown to be safe to use for individuals with dementia, specifically reducing agitation and increasing social contact in this population. [9,10] Based on our preclinical study, we present MLD as a potential therapy that may improve cognition and disease-modifying biomarkers in AD. Moreover, as MLD is already approved in humans, it can be translated immediately as a stand-alone or adjunct therapy.

## CRediT Statements

Mitchell J. Bartlett: Conceptualization, Methodology, Validation, Formal Analysis, Investigation, Resources, Data Curation, Writing – Original Draft, Writing – Review & Editing, Visualization, Supervision, Project Administration, Funding Acquisition.

Robert P. Erickson: Conceptualization, Writing – Review & Editing.

Jennifer B. Frye: Validation, Investigation, Resources, Writing – Review & Editing.

Kristian P. Doyle: Validation, Resources, Writing – Review & Editing, Supervision.

Paulo W. Pires: Conceptualization, Methodology, Validation, Formal Analysis, Investigation, Resources, Data Curation, Writing – Review & Editing, Visualization, Supervision, Project Administration, Funding Acquisition.

Marlys H. Witte: Conceptualization, Resources, Writing – Review & Editing, Funding Acquisition.

## Acknowledgements

We would like to thank Michael J. Bernas for his guidance and suggestions in adapting the manual lymph drainage procedure to mice. The mouse strain used for this research project, B6SJL-Tg (APPSwFILon, PSEN1*M146L*L286V)6799Vas/Mmjax, RRID:MMRRC 0384840-JAX, was obtained from the Mutant Mouse Resource and Research Center (MMRRC) at The Jackson Laboratory, a National Institutes of Health (NIH)-funded strain repository, and was donated to the MMRRC by Robert Vassar, PhD, Northwestern University. This work was supported by grants from the Arizona Alzheimer’s Consortium, Arizona Department of Health Services to Mitchell J. Bartlett, Paulo W. Pires, and Marlys H. Witte. Additional funding was provided by the Van Winkle Fund and the NIH NINDS (R25 NS 076437) to Marlys H. Witte.

